# Characterization and hierarchy of the spermatogonial stem cell compartment in human spermatogenesis by spectral cytometry using a 16-colors panel

**DOI:** 10.1101/2023.12.21.572912

**Authors:** C. Lapoujade, M. Blanco, M. Givelet, A.S Gille, I. Allemand, L. Lenez, N. Thiounn, S. Roux, J.P. Wolf, C. Patrat, L. Riou, V. Barraud-Lange, P. Fouchet

## Abstract

About one in six couples experience fertility problems, and male infertility accounts for about half of these cases. Spermatogenesis originates from a small pool of spermatogonial stem cells (SSCs), which are of interest for the treatment of infertility but remain poorly characterized in humans. Using multiparametric spectral flow cytometric analysis with a 16-colours (16-C) panel of cell markers, we identify novel markers of SSCs and provide insights into unravelling and resolving the heterogeneity of the human spermatogonial cells. This 16-C panel of markers allowed the identification of a primitive SSCs state with the β-2M^−^CD51/61^−^ ITGA6^+^SSEA4^+^TSPAN33^+^THY1^+^CD9^+^EPCAM^med^CD155^+^CD148^+^CD47^high^CD7^high^ phenotype, with a profile close to the most primitive SSCs states 0 and SSC1-B previously defined by sc-RNAseq approach. The hierarchy of events in the spermatogonial stem cell and progenitor compartment of human spermatogenesis has been delineated. This highlights the importance of a multi-parametric and spectral cytometry approach. The in-depth characterization of testicular cells should help to overcome the lack of stem cell knowledge, that hinders the understanding of the regenerative potential of SSCs, and is a critical parameter for the successful development of new SSCs-based cell therapies.

## INTRODUCTION

Regenerative medicines based on spermatogonial stem cells (SSCs) are of great interest to treat for infertility issues. Efficient production of functional sperm after testicular transplantation has been reported in non-human primates, as well as successful autografting of testicular tissue to produce sperm ^1,2^. As in other tissue undergoing continuous renewal, the pool of SSCs ensures adult spermatogenesis by self-renewing and differentiating to produce sperm. Basic knowledge of SSCs has benefited greatly from studies in Caenorhabditis elegans, Drosophila and mouse models. SSCs identity based on a hierarchical pattern or a more stochastic model tuning the differentiation of a larger subset of SSCs and early progenitors equipotent in term of stemness, emerged to regulate the fate of the stem cell pool ^3–5^.

In human, the spermatogonial stem cell pool and the molecular mechanisms governing their fate are still poorly understood, although recent scRNAseq studies have made remarkable contributions to define different cell SSCs states in the primitive spermatogonial pool ^6–11^. However, such an approach is also limited by the inability to isolate these cell subsets to study their function. The multidimentional approach of scRNAseq highlights the need to use multiple cellular parameters simultaneously in single cells to characterize the heterogeneity of the spermatogonial pool with high resolution. The flow cytometry approach and recent developments such as spectral cytometry, which allow high-throughput multi-parametric phenotyping of single cells in suspension, could facilitate the characterization of cell phenotype and functionality, as well as the resolution of the complexity of the human SSCs pool necessary for the development of new therapies.

Here, we have generated a 16-colour (16-C) panel of markers that allows high-throughput multi-parametric and in-depth phenotyping of single testicular cells using spectral cytometry, facilitating the characterization of the different germinal populations. We identified novel markers of SSCs and provided insights to unveil and resolve the heterogeneity of the human spermatogonial cell heterogeneity in a non-deleterious manner for cell integrity and viability, allowing future studies of their functionality.

## RESULTS

### Development of a 16-C panel to characterize human spermatogenesis and discriminate SSCs populations using spectral flow cytometry and high-throughput multi-parametric analysis

We recently identified a spermatogonial subpopulation highly enriched in SSCs with the phenotype Side population(SP)β2M^−^ITGA6^+^THY1^+^ population using a 5-color panel ^12^. Based on this set of markers, we aimed to increase the panel to apply a multi-parameter flow cytometry strategy to further characterize human spermatogenesis and improve the delineation of SSCs identity. First, TSPAN33, FGFR3 and SSEA4, previously described to be expressed on SSCs and primitive spermatogonia, were added to our β2M, ITGA6, and THY1 panel of markers, along with the KIT receptor allowing to discriminate differentiating from primitive spermatogonia ^6,9,12–14^. The β2M^−^ITGA6^+^THY1^+^ population contains the spermatogonia expressing SSEA4, FGFR3, and the fraction of cells expressing TSPAN33 (Figure S1A and B). TSPAN33 expression in the SSEA4+ population clearly discriminated a subpopulation representing 0.26% of germ cells that co-expressed THY1 but were negative for the KIT receptor (Figure S1C). To enrich this 8-C panel with new markers, we screened 17 membrane receptors previously described in transcriptomes or sc-RNAseq from human germ cell populations, or markers of murine SSCs, or markers of stem cells from other tissues ^6–11,14^. Among these, a fraction of β2M^−^ITGA6^+^THY1^+^ cells were positive for the expression of CD47, CD155, CD7, CD130 or CD148 (Figure S2). Finally, we identified meiotic and post-meiotic populations according to the expression of CD148, CD155, CD47 and ITGA6 (Figure S3).

To further extend the multi-parametric and in-depth characterization of human SSCs and spermatogenesis, we combined all these markers and profiled adult testicular cell samples by spectral flow cytometry using 15 antibodies and a viability marker (β-2M, CD51/61, ITGA6, SSEA4, TSPAN33, FGFR3, THY1, CD9, EPCAM, CD47, CD7, CD148, CD155, CD130, KIT, DAPI/viability) resulting in a 16-C panel. The different signals were resolved by spectral deconvolution and the autofluorescence of the cells was extracted (Figure 1 and Figure S4), allowing the analysis of 18 parameters (16 markers and FSC/SSC). Figure 1 shows the manual gating strategy for the 16-color panel used in this study, and representative flow cytometry plots of the 13 germinal markers displaying every parameter versus every other parameter are presented in Figure S4. First, single living (DAPI-negative) testicular cells were selected and spermatozoa were removed from analysis by gating based on FSC and SSCs from 405 nm laser. Testicular cells were then divided according to somatic cell marker β2M, and the integrin complex CD51/CD61 known as α_V_β_3_ CD51/CD61 is expressed in testicular Leydig stem cells and macrophages, and is used in the murine marker panel to negatively discriminate spermatogonia ^15–17^. As shown in Figure 1A, human testicular CD51/61^+^ cells are also positive for β2-microglobulin. Cells negative for CD51/61 and β2M markers were then gated for germ cell analysis. SSEA4^+^ cells expressed ITGA6 and CD9. The expression of ITGA6, THY1, CD47, TSPAN33, CD7, FGFR3, CD155 and SSEA4 was detected in EPCAM^med^ cells (Figure 1A and Figure S4), intermediate expression of EPCAM being a marker of primitive spermatogonia ^7^. SSEA4^+^ cells were positive for the CD148 marker, and the majority did not express the KIT receptor, although KIT was found in cells with medium expression of SSEA4. We then subdivided the spermatogonial populations using SSEA4 and TSPAN33 markers (Figure 1A). The other markers were analysed according to the defined SSEA4^+^TSPAN33^+^ (0.09 % of the germ population-Figure 1B), SSEA4^+^TSPAN33^−^ (1.1% of the germ population-Figure 1C) and SSEA4^−^ (Figure 1D) cell populations. We observed EPCAM^med^ cells with higher expression levels of CD7, CD47, THY1, CD148, FGFR3 and CD155 in the SSEA4^+^TSPAN33^+^ population when compared to SSEA4^+^TSPAN33^−^ cells.

**Figure 1:**
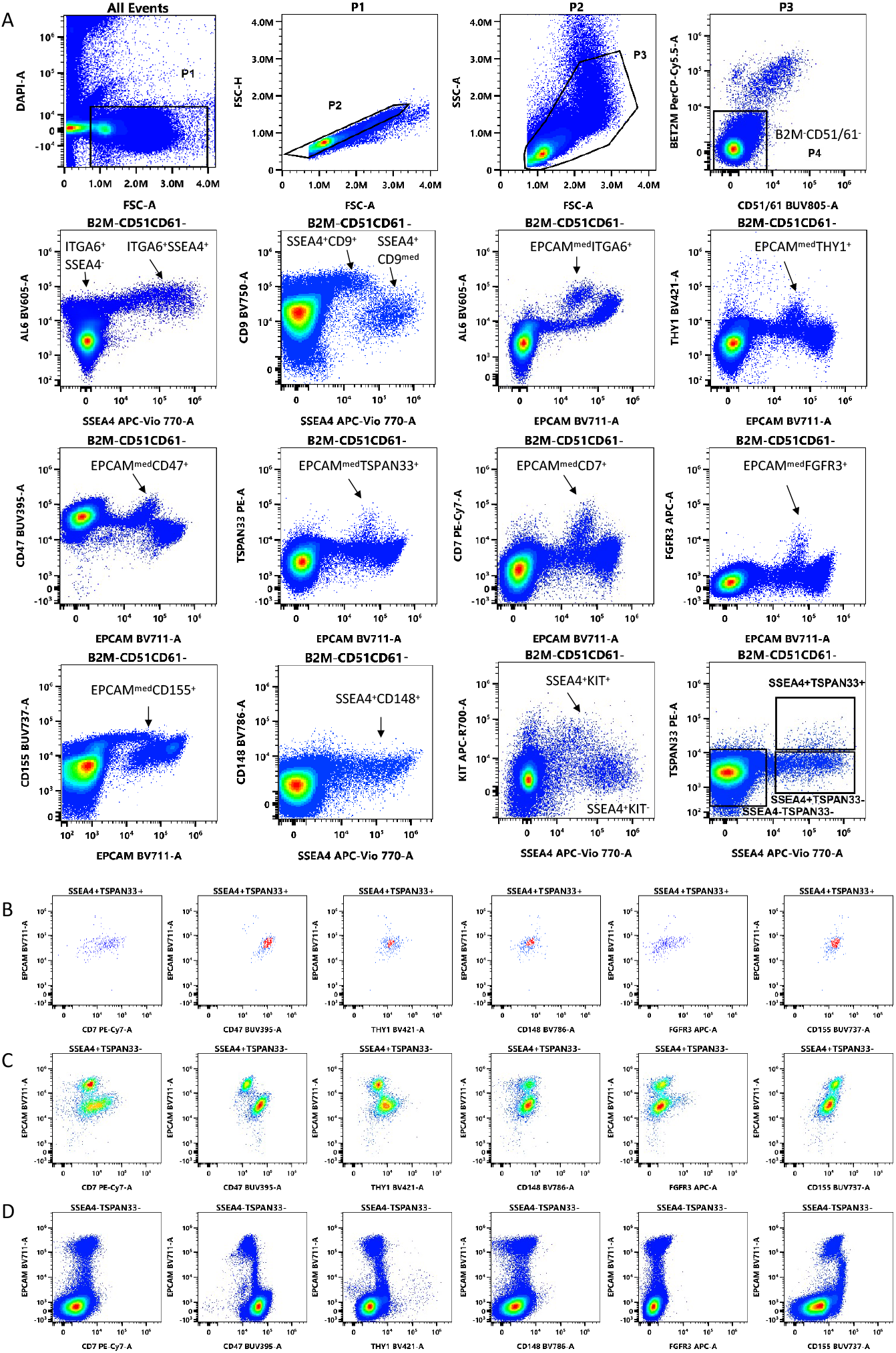
Flow cytometry profiling of adult testicular cell samples using the 16-C panel. (A) Representative flow cytometry plots of the spectral flow analysis on the human spermatogonial progenitors according to FSC, SSC, β2M, CD51/CD61, ITGA6, THY1, SSEA4, and TSPAN33 parameters. Testicular cells were gated based on morphology to remove cellular debris, sperm and doublets. (B, C, D) Expression of CD7, CD155, CD148, CD47, and KIT markers in the β2M/CD51/CD61^−^SSEA4^+^TSPAN33^+^ (B), the β2M/CD51/CD61^−^SSEA4^+^TSPAN33^−^ (C), and the β2M/CD51/CD61^−^SSEA4^−^TSPAN33^−^ (D) germinal populations. Figure S5 shows the most relevant fluorescence minus one (FMO) controls. ITGA6 is noted AL6 in the cytogram axis.

### Comprehensive analysis of the multidimensional 16-C panel dataset of human testicular cells

To allow comprehensive analysis of the resulting multi-dimensional dataset, the total viable testis cells (341 378 cells) were first projected into the same UMAP space (Figure 2A). The spermatogonial, meiotic and postmeiotic cell subsets obtained by manual gating, as shown in Figure 1, Figure S3 and Figure S4, were overlaid on the UMAP dimensionality reduction plot (Figure 2B). All cell subsets formed well-defined clusters. We discriminated somatic cell populations (β2M^+^CD51/61^+^and β2M^+^CD51/61^−^ cells) from germ populations. We distinguished meiotic spermatocytes I (4N), and spermatocytes II (2N) and postmeiotic (N) cells from spermatogonial populations. In addition, primitive spermatogonial populations (ITGA6^+^THY1^+^ and ITGA6^+^SSEA4^+^ cells) were clearly separated from differentiating spermatogonia, demonstrating that the dimensionality reduction visualization was able to separate minor populations of spermatogonia. Heatmap overlay of all the markers on the UMAP plot shows that SSEA4, ITGA6, THY1, CD7, TSPAN33, CD47, EPCAM^dim^, CD9, FGFR3, CD148 and CD155 markers, but not KIT, were expressed in the primitive spermatogonial ITGA6^+^THY1^+^ subset (Figure 2C). We also observed a continuum of differentiation from the primitive and differentiating spermatogonial clusters to spermatocytes I (4N), spermatocytes II (2N) and spermatid cells (Figure 2A and B). Unsupervised clustering of β2M^+^CD51/61^−^ germ cells made by FlowSOM confirmed our manual clustering of germ cells (Figure S6). A FlowSOM tree and a trajectory inference were derived from the unsupervised clustering of the β2M^−^ CD51/61^−^ germinal cells. The FlowSOM tree recapitulated the successive steps of the differentiation process (Figure 3A and Figure S6). The resulting heatmap from the trajectory inference shows the expression levels of the different markers along this trajectory (Figure 3B). Based on the expression of markers (Figure S6), the germline clusters were overlaid on the trajectory (Figure 3B). This confirmed that the trajectory inferred from the unsupervised analysis of the 16-C panels was successful in describing the spermatogonial differentiation.

**Figure 2.**
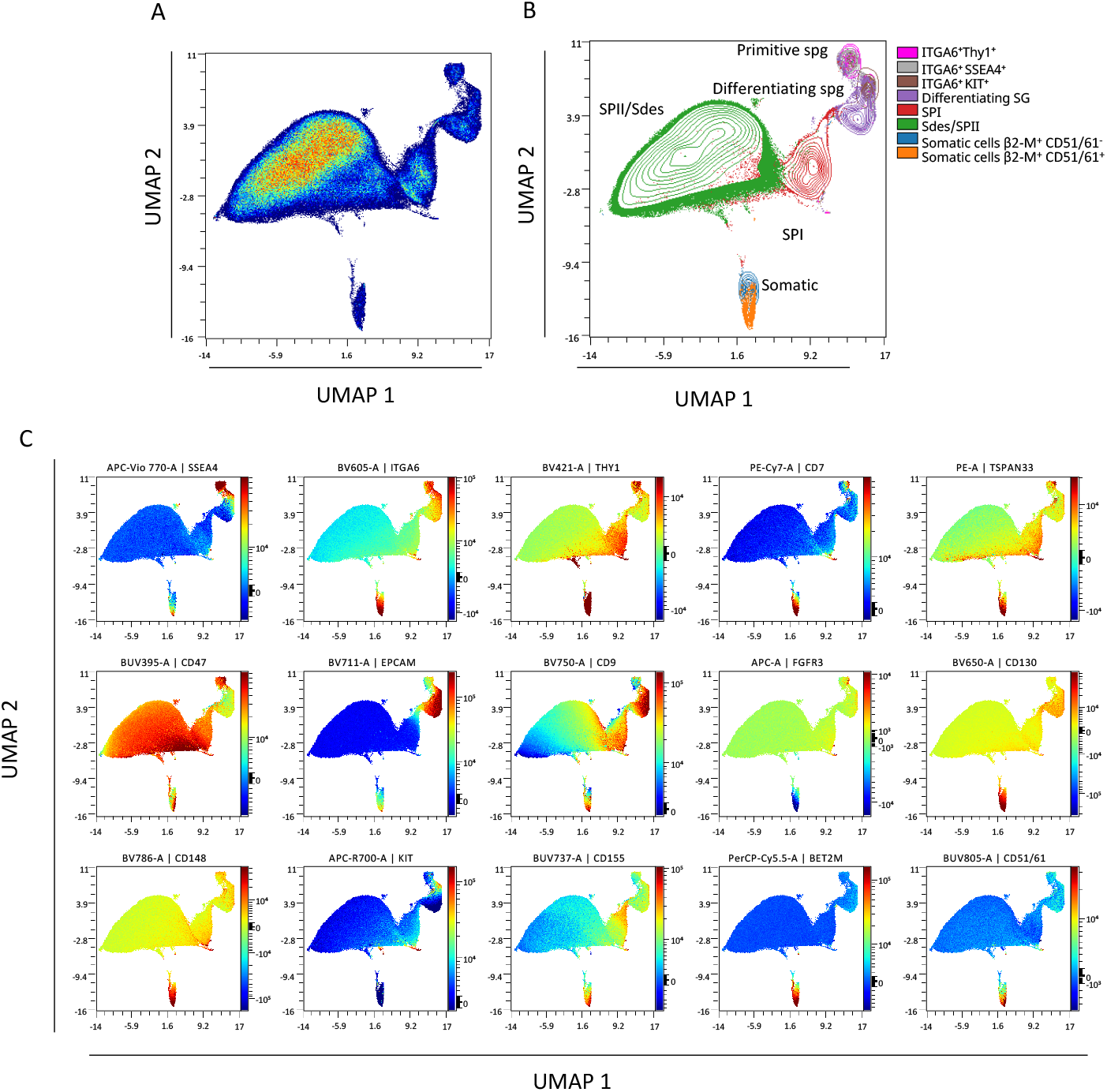
Comprehensive analysis of multidimensional 16-C panel dataset of human testicular cells. (A) High-dimensional UMAP analysis of human testicular cells, (B) Testicular cell subsets defined in Fig. 1 and Fig. S4 are indicated on the UMAP. Spermatocyte I (SPI), spermatocyte II (SPII), spermatids (Sd), and the “S phase” subpopulation (differentiating spermatogonia) were assigned according to the expression of CD148, CD47, CD155 and ITGA6 as previously defined in Figure S3 and S4. (C) Expression patterns of individual spermatogonial markers overlaid onto the UMAP representation of human testis cells. The colour indicates the fluorescence intensity of the marker (red: high to blue: neg).

**Figure 3.**
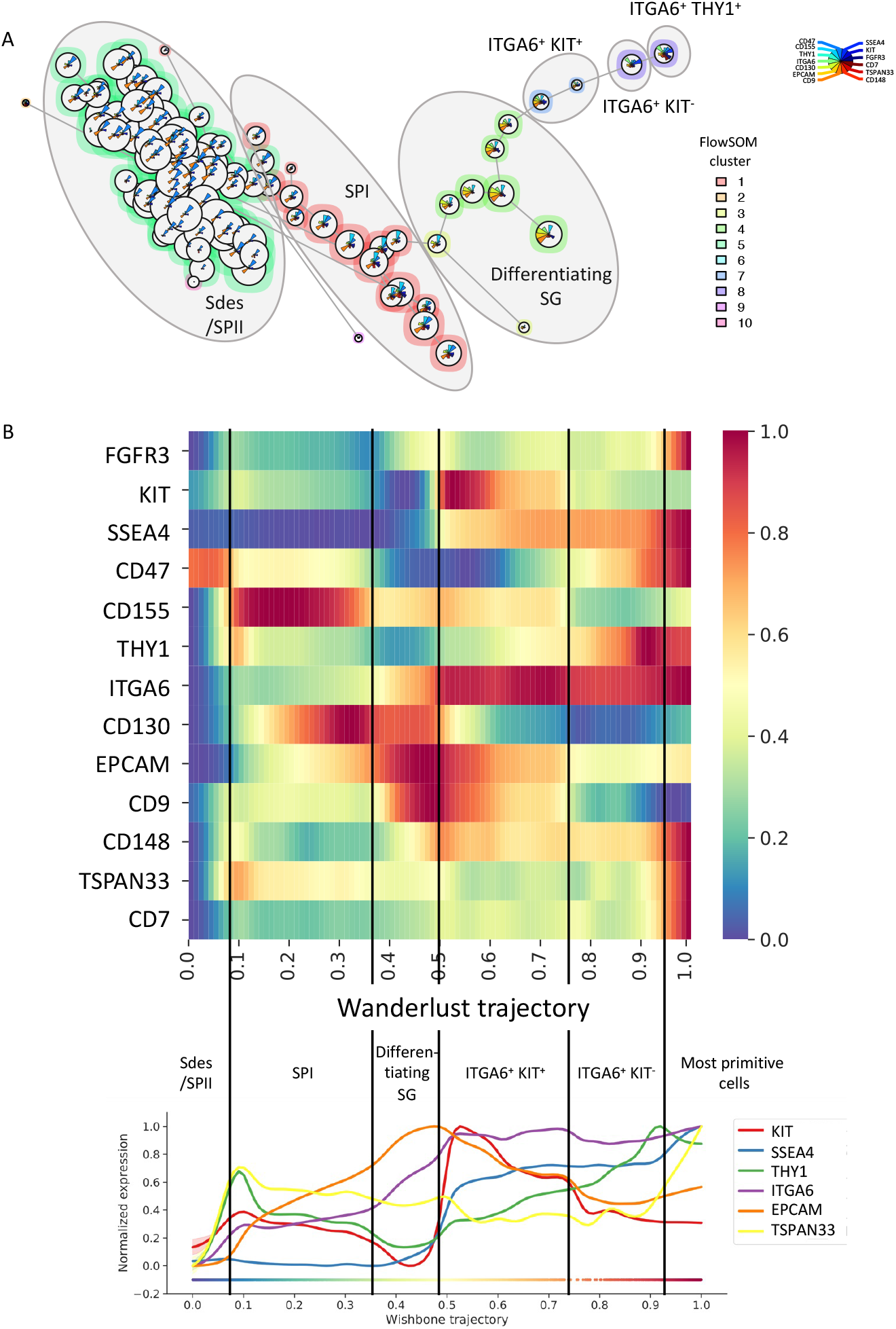
Trajectory analysis of the human germ cell differentiation dataset (A) FlowSOM tree inferred from unsupervised clustering of β2M^+^CD51/61^−^ germinal cells using FlowSOM analysis. Germinal subsets are indicated on the FlowSOM tree plot. (B) Heatmap and Wanderlust trace of the expression of the germinal markers after Wanderlust trajectory inference. The different differentiation stages are indicated on the heatmap and Wanderlust trace to show the expression patterns of each differentiation state.

### Delineation of the human spermatogonial progenitors and stem cells populations using the 16-C panel

We then focused the analysis on the whole β2M^−^CD51/61^−^ITGA6^+^ spermatogonial population. Adult testicular cell suspensions from 3 patients with normal spermatogenesis were phenotyped and the data were concatenated to include a higher number of spermatogonia in the analysis. 141 451 β2M^−^CD51/61^−^ITGA6^+^ were projected into UMAP spaces (Figure 4A), and the major spermatogonial subsets obtained by manual gating were overlaid on this UMAP plot (Figures 4B). Three main cell populations were delineated (1) a primitive spermatogonial cluster consisting of ITGA6^+^SSEA4^+^ and ITGA6^+^THY1^+^ populations, with SSEA4^+^TSPAN33^+^ cells at the edge of this cluster, (2) the ITGA6^+^KIT^+^ differentiating spermatogonial cluster making a cell state transition, (3) towards the third cluster of more advanced differentiated ITGA6^+^SSEA4^−^ spermatogonia. Unsupervised FlowSOM analysis allowed to distinguish 6 main clusters (clusters 1, 5, 6, 7, 8 and 10) according to the expression of the 13 markers (Figure S7), allowing to distinguish primitive spermatogonia (clusters K6, K8 and K10) from more advanced spermatogonial progenitors (clusters K1, K5 and K7). Heatmap overlay of all the 13 markers on UMAP plot showed a differential expression of CD47, TSPAN33, CD7, THY1, SSEA4, ITGA6, EPCAM, FGFR3, CD148, CD9 and CD155 markers in the primitive spermatogonial cluster (Figures 4C and D), allowing the discrimination of different subpopulations in this cluster, with in particular, at its lower edge a prominent expression for CD47, TSPAN33, CD7, ITGA6, FGFR3, CD148, CD9 and CD155, and a medium expression of EPCAM. Higher expression of EPCAM, CD148, CD9, and CD130 was also observed in the more advanced differentiated ITGA6^+^KIT^+^ and ITGA6^+^SSEA4^−^ spermatogonia, together with a lower ITGA6 expression.

**Figure 4:**
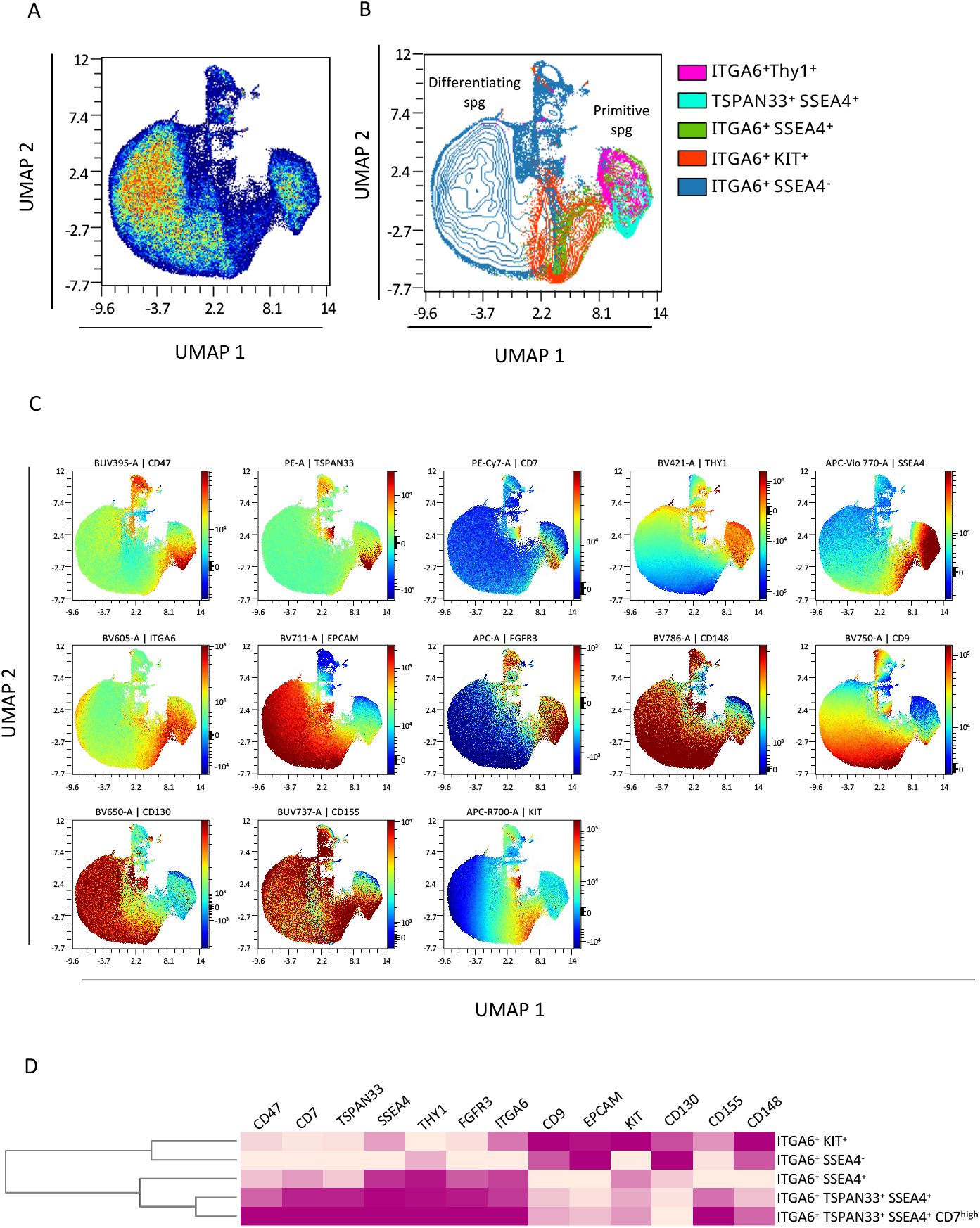
Comprehensive analysis of multidimensional 16-C panel dataset of ITGA6^+^ spermatogonia. (A) High-dimensional UMAP analysis of human ITGA6^+^ spermatogonia, (B) Testicular cell subsets defined in Fig. 1, Fig. S3 and Fig. S4 are indicated on the UMAP. (C) Expression patterns of individual spermatogonial markers overlaid onto the UMAP. Colour indicates fluorescence intensity. (D) Heatmap of differentially expressed markers in the spermatogonial populations

### In-depth characterization of the human β2M^−^CD51/61^−^ITGA6^+^THY1^+^ SSCs population

When 9074 β2M^−^CD51/61^−^ITGA6^+^THY1^+^ primitive spermatogonia from 3 patients with normal spermatogenesis were projected on UMAP (Figure 5), we observed mainly two clusters, one consisting of SSEA4^+^TSPAN33^+^, SSEA4^+^TSPAN33^+^CD47^high^, and SSEA4^+^ TSPAN33^+^CD7^high^ cells on the left side, and one consisting of SSEA4^+^TSPAN33^−^ cells and ITGA6^+^CD155^med^ on the right side (Figures 5A, B and C). Heat maps of marker expression shows a progressive expression of the CD7 marker from high to low levels as one progresses from SSEA4^+^TSPAN33^+^ cluster to SSEA4^+^TSPAN33^−^ one (Figure 5D). SSEA4^+^TSPAN33^+^ cells show the higher expression of CD47, CD7, CD9, and CD155 markers and intermediate expression of CD130 and CD148 (Figure 5D). FGFR3 expression was found in cells at the border of the two clusters, suggesting a transitional step between the two states. Unsupervised clustering of β2M^−^CD51/61^−^ITGA6^+^THY1^+^ cells by FlowSOM mainly identified three clusters K1, K3 and K4 (Figure S8B), SSEA4^+^TSPAN33^−^ population being divided in two clusters K1 and K4. The expression profile of K3 corresponded to an immature phenotype (expression of TSPAN33, CD47, CD7, EPCAM, THY1, and FGFR3), K1 to a more mature phenotype (lower expression of TSPAN33, CD47, CD7, SSEA4, THY1, FGFR3, ITGA6, EPCAM, CD155, CD148, and higher expression of KIT and CD130), K4 to an intermediate phenotype (Figure S8C). Surprisingly, the K1 cluster was placed between the K3 and K4 clusters in the FlowSOM tree, although cells from the K1 cluster display the most mature phenotype (Figure S8D). This finding deviates from the expected linear process starting from K3 going to K4, and then to K1 cells. We postulate that K3 and K4 cells could represent alternative stem cell states with different potential, with K3 being the least developmentally advanced SSCs state. K3 and K4 states would converge on the K1 state to commit to differentiation. The existence of different subsets of SCCs that may interconvert between them or have different potentials has yet been described^6,9^. Finally, we investigated whether the population we defined as SSEA4^+^TSPAN33^+^ could correspond to the state 0 of the spermatogonial hierarchy previously defined in scRNAseq analysis ^6^. Consistent with this, we found that mRNA of *C19orf84, PIWIL4, FGFR3, UTF1*, and *LPPR3* markers of state 0 and state SSC1-B ^6,9^ were highly expressed in β2M^−^ SSEA4^+^TSPAN33^+^ cell fractions compared to β2M^−^SSEA4^+^TSPAN33^−^ cells (Figure 5E), which were flow sorted using the 8-C panel and ARIA flow cytometer (see Figure S1). Therefore, the β-2M^−^CD51/61^−^ITGA6^+^SSEA4^+^TSPAN33^+^THY1^+^CD9^+^EPCAM^med^CD155^+^CD148^+^CD47^high^CD7^high^cell population should correspond to the most primitive SSCs state 0 or state SSC1-B, or include these primitive states of SSCs ^6,9^.

**Figure 5:**
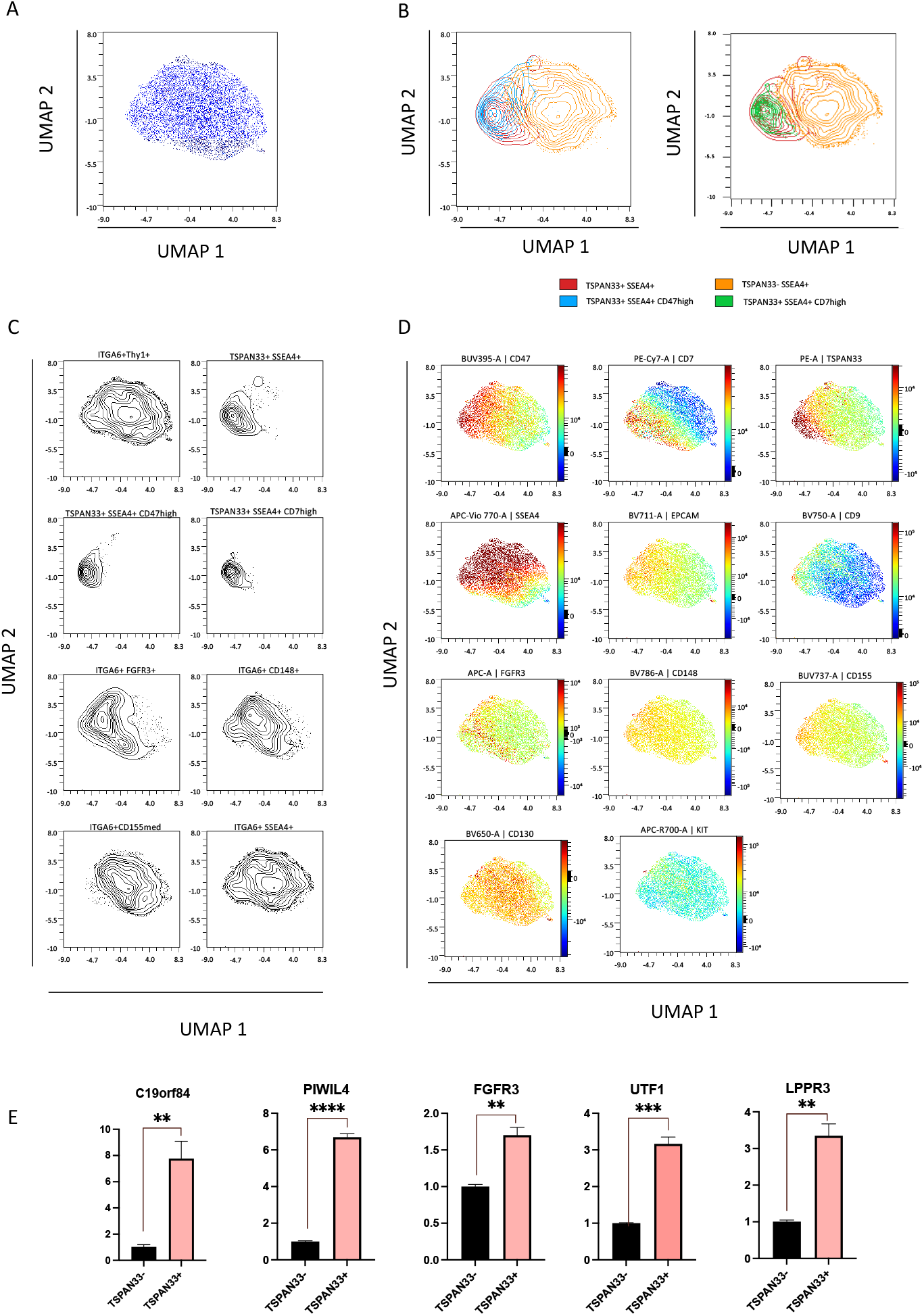
Comprehensive analysis of multidimensional 16-C panel dataset on the ITGA6^+^THY1^+^ spermatogonial population. (A) High-dimensional UMAP analysis of β2M/CD51/CD61^−^ITGA6^+^THY1^+^ cells (UMAP zoom to main cluster, full UMAP is shown in Figure S8A). (B) Overlay of SSEA4^+^TSPAN33^+^, SSEA4^+^TSPAN33^−^, SSEA4^+^TSPAN33^+^CD47^high^ and SSEA4^+^TSPAN33^+^CD7^high^ populations as defined in Fig. 1 and Fig. S4. (C) Spermatogonial cell subsets defined in Fig. 1 and Fig. S4 are shown on the UMAP. (D) Expression patterns of individual spermatogonial markers overlaid onto the UMAP. Colour indicates fluorescence intensity (red: high to blue: neg) (E) Expression of *C19orf84, PIWIL4, FGFR3, UTF1* and *LPPR3* markers in sorted β2M^−^SSEA4^+^TSPAN33^+^ and β2M^−^SSEA4^+^TSPAN33^−^ populations (n=3).

## DISCUSSION

The understanding of the developmental hierarchy of human SSCs and spermatogonial progenitors has greatly benefited from scRNAq studies, allowing the delineation of different cell states in the primitive spermatogonial population ^6–11^. However, the functional studies of these putative SSCs subsets are still limited. The TSPAN33 and LPPR3 markers identified in these subsets were recently confirmed to define a cell population with a regenerative potential using testicular transplantation assay ^10,14^. Here, we designed a 16-C panel to generate a comprehensive profile of human SSCs by spectral flow cytometry. A comprehensive screening of extracellular membrane markers of SSCs, embryonic stem cells, and other types of tissue stem cells allowed us to identify novel markers of SSCs such as CD47, CD7, CD148, and CD155. This combination allows the identification of a primitive SSCs state with β-2M^−^CD51/61^−^ITGA6^+^SSEA4^+^TSPAN33^+^THY1^+^CD9^+^EPCAM^med^CD155^+^CD148^+^CD47^high^CD7^high^ phenotype.

Profiling the SSCs subsets suggests a hierarchy in the differentiation process with a transition from a primitive ITGA6^+^SSEA4^+^TSPAN33^+^THY1^+^CD9^+^EPCAM^med^CD155^+^CD148^+^ CD47^high^CD7^high^ state to an ITGA6^+^SSEA4^+^TSPAN33^−^THY1^+^CD9^+^EPCAM^med^CD130^+^CD155^med^ CD148^+^CD47^+^CD7^low/med^ state, FGRG3^+^ cells are found at the border of the 2 cell clusters suggesting a transition step. They then differentiate into ITGA6^+^SSEA4^+^TSPAN33^−^CD7^−^THY1^−^ CD9^+^EPCAM^+^CD130^+^CD155^+^CD148^+^ KIT^+^ differentiating spermatogonial states. The expression of KIT represents a cell state transition towards the third cluster of more advanced differentiated ITGA6^+^SSEA4^−^EPCAM^+^CD130^+^CD9^+^CD155^med^ spermatogonia. Unsupervised FlowSOM clustering of β2M^−^CD51/61^−^ITGA6^+^THY1^+^ primitive spermatogonia suggests that this population is composed of three SSCs clusters, two of which (K1 and K4) are putative alternative stem cell states, both linked to the K3 cluster, which represents the least developmentally advanced SSC state. The potential interconversion between different SSCs substates has already been postulated in humans ^6,9^, and is reminiscent of the alternative non-linear mouse stem cell model ^4^.

TSPAN33, SSEA4, ITGA6, THY1, and LPPR3 are markers of spermatogonial cells with repopulation potential ^10,12–14^. Notably, we show that β2M^−^CD51/61^−^SSEA4^+^TSPAN33^+^ cells coexpress the stem cell markers THY1 and LPPR3. In addition, TSPAN33^+^ SSCs express CD47 at higher levels compared to the TSPAN33^−^ population. CD47 expression has been described to be enriched in the transcriptome of LPPR3^+^ and SPβ2M^−^ITGA6^+^THY1^+^ SSCs population that exhibit regenerative potential after transplantation ^12,14^. CD47 is also a marker of state 0 and state SSC1-B, which are considered to be the most primitive SSCs ^6,9^. The coexpression of TSPAN33 and CD47 expression, together with *PIWIL4, C19orf84, LPPR3* and *UTF1* in this subpopulation of SSCs, representing 0.13% of the germinal population, suggests that it corresponds to the primitive primitive state 0 and SSC1-B of SSCs, previously described in scRNAseq studies^6,9,10,14^.

We show that CD47, CD155, CD148, and CD7 are expressed in the populations of human primitive spermatogonia, and question their potential role in the physiology of SSCs. The main role of the transmembrane protein CD47 is to protect cells from phagocytosis. CD47 when bound to regulatory protein-α (SIRPα) expressed on macrophages and dendritic cells, inhibits the process of phagocytosis ^18^. For example, mobilised haematopoietic stem cells protect themselves from phagocytosis by modulating CD47 expression ^19^. As testicular resident macrophages have been described to take place in the niche layered at the basement membrane, higher CD47 expression may be a way to protect SSCs from potential phagocytosis by macrophages. Recently, a role for SIRPα/CD47 signaling was also suggested in selfrenewal of murine SSCs ^20^. CD155, originally identified as poliovirus receptor (PVR) and expressed in undifferentiated spermatogonia from prepubertal mouse testes^21^, is a member of the immunoglobulin superfamily involved in many different physiological processes mediating intercellular interactions or cell adhesion to the extracellular matrix, and cell migration^22^. CD148 (PTPRJ), a receptor-type protein tyrosine phosphatase, has a prominent role in the negative regulation of growth factor signalling and cell proliferation ^23,24^. In particular, it has been reported to modulate the FGF receptor (FGFR), Erk1/2, and the the Src family of protein tyrosine kinases (SFKs) ^25^, molecular pathways that have been shown to be involved in the regulation of murine SSCs fate ^26,27^. CD7 is a T-cell associated molecule expressed early in T cell ontogeny and is a marker of various cancer stem cells, but the role of CD7 remains poorly understood beyond its role in T cell activation ^28^.

The development of mass cytometry and spectral cytometry has expanded the ability to more accurately define cell heterogeneity in complex biological systems, allowing a large number of cell phenotyping data to be collected simultaneously on single cells. Here, we have characterized the dynamic changes in the expression of markers of human SSCs using spectral flow cytometry. The combination of a larger number of biomarkers and computational approaches should help to reduce the knowledge gap in the identification of human primitive spermatogonia, allowing an accurate characterization of the spermatogonial populations and a better ranking of the successive SSCs states by analysing their functionality.

## Experimental procedures

### Human materials

Adult human testis biopsies from obstructive azoospermia patients with normal spermatogenesis (32, 37 and 45 years old for 16-C panel analysis) were obtained from the CECOS Hospital Cochin. All patients consented to inclusion in this research study (IRB-approved protocol: IRB 00003835; 2012/40ICB; French Institutional Review Board-Comité de Protection des Personnes, Ile de France IV).

### Testicular single-cell suspensions, immunomagnetic and flow cell sorting, and analysis

Testicular single-cell suspensions were prepared from human biopsies. The tissue was incubated in trypsin 0.25% containing collagenase I (final concentration, 0.5 mg/ml) for 20 minutes at 34°C. The cell suspension was then filtered (20 μm). The mix of antibodies was incubated with Brilliant Stain Buffer Plus (BD Biosciences) and cells were preincubated in Human TruStain FcX™(Biolegend) and True-Stain Monocyte Blocker™ (Biolegend) according to manufacturer instruction. Then cells were incubated with the following antibodies for 20 min at 4°C as previously described ^29^ : anti-ITGA6, anti-β2-M, anti-CD148, anti-CD155, anti CD51/61, anti-CD47, anti-CD9, anti-CD130, anti-KIT (all from BD Biosciences), anti-TSPAN33, anti-THY1, anti-EPCAM and anti-CD7 (all from Biolegend), anti-SSEA4 (Miltenyi Biotec), and anti-FGFR3 (Biotechne). After washing, propidium iodide (Sigma), or DAPI (Sigma) was added before cell sorting to exclude dead cells. Analysis and cell sorting were performed using FACSAria cytometer (BD Biosciences) and spectral 5 lasers Aurora cytometer (Cytek). Spectral flow cytometer was calibrated using standard beads following provider specifications. All reference controls, except DAPI, were acquired using 1 drop UltraComp eBeads compensation beads (Invitrogen) mixed with antibody for reference controls. Spill over and gate positions were determined using fluorescence minus one (FMO) controls to delineate boundaries separating negative from positive staining. SpectroFlo software (Cytek Biosciences), which uses Ordinary Least Squares Linear Unmixing was used to deconvolute the different fluorescence spectra ^30^. The data were analyzed with DIVA, FlowJo, or SpectoFlo software. UMAP analysis were performed on indicated populations using OMIQ (Dotmatics) as an unsupervised nonlinear dimensionality reduction method to explore and visualize the multidimensional data generated with the panel based on all parameters (excluding forward scatter and side scatter). FLowSOM ^31^ and trajectory analysis was done using the cytometry analysis platform OMIQ (Dotmatics). Wanderlust analysis ^32^ was realized using diffusion maps as a dimensionality reduction algorithm, the trajectory is inferred using the eigenvector (variance in the data) of the diffusion component, and the different testicular markers.

### RNA extraction and quantitative RT-PCR

mRNA was prepared using RNeasy® Micro and Mini kits (Qiagen). The mRNA was then reverse-transcribed with a Quantitect kit (Qiagen). Quantitative RT-PCR was performed using an AB7900 device (Applied Biosystems) with Fast SYBR® Green Master Mix (Applied Biosystems). The primers are listed in Table S1.

### Statistics

All values are shown as the means ± SEM. Statistical analyses were performed by Student’s t test (GraphPad Prism software): ns, not significant, p > 0.05; *, p < 0.05; **, p < 0.01; ***, p < 0.001.

## ACKNOWLEDGEMENT

The authors acknowledge the support of the members of the CECOS from CHU Cochin. This work was supported by grants from INCA, the Agence de Biomédecine, the “Ligue contre le cancer”, and EDF.

## AUTHOR CONTRIBUTIONS

CL, MB, VBL and PF provided project design. CL, MB, MG, LR and PF performed experiments, analyzed and interpreted data. ASG, IA, LL, AJ, FH, IA and LR participated in experimentation and provided comments. ASG, NT, SR, PW, CP, and VBL supervised the sample acquisition, CL and PF prepared the manuscript.

## REFERENCES

1. Hermann, B.P., Sukhwani, M., Winkler, F., Pascarella, J.N., Peters, K.A., Sheng, Y., Valli, H., Rodriguez, M., Ezzelarab, M., Dargo, G., et al. (2012). Spermatogonial stem cell transplantation into Rhesus testes regenerates spermatogenesis producing functional sperm. Cell Stem Cell 11, 715–726. 10.1016/j.stem.2012.07.017.

2. Fayomi, A.P., Peters, K., Sukhwani, M., Valli-Pulaski, H., Shetty, G., Meistrich, M.L., Houser, L., Robertson, N., Roberts, V., Ramsey, C., et al. (2019). Autologous Grafting of Cryopreserved Prepubertal Rhesus Testis Produces Sperm and Offspring. Science 363, 1314–1319. 10.1126/science.aav2914.

3. Oatley, J.M., and Brinster, R.L. (2012). The germinal stem cell niche unit in mammalian testes. Physiol Rev 92, 577–595. 10.1152/physrev.00025.2011.

4. Hara, K., Nakagawa, T., Enomoto, H., Suzuki, M., Yamamoto, M., Simons, B.D., and Yoshida, S. (2014). Mouse Spermatogenic Stem Cells Continually Interconvert between Equipotent Singly Isolated and Syncytial States. Cell Stem Cell 14, 658–672. 10.1016/j.stem.2014.01.019.

5. Rooij, D.G. de (2017). The nature and dynamics of spermatogonial stem cells. Development 144, 3022–3030. 10.1242/dev.146571.

6. Guo, J., Grow, E.J., Mlcochova, H., Maher, G.J., Lindskog, C., Nie, X., Guo, Y., Takei, Y., Yun, J., Cai, L., et al. (2018). The adult human testis transcriptional cell atlas. Cell Res 28, 1141–1157. 10.1038/s41422-018-0099-2.

7. Hermann, B.P., Cheng, K., Singh, A., Roa-De La Cruz, L., Mutoji, K.N., Chen, I.-C., Gildersleeve, H., Lehle, J.D., Mayo, M., Westemströer, B., et al. (2018). The Mammalian Spermatogenesis Single-Cell Transcriptome, from Spermatogonial Stem Cells to Spermatids. Cell Rep 25, 1650-1667.e8. 10.1016/j.celrep.2018.10.026.

8. Wang, M., Liu, X., Chang, G., Chen, Y., An, G., Yan, L., Gao, S., Xu, Y., Cui, Y., Dong, J., et al. (2018). Single-Cell RNA Sequencing Analysis Reveals Sequential Cell Fate Transition during Human Spermatogenesis. Cell Stem Cell 23, 599-614.e4. 10.1016/j.stem.2018.08.007.

9. Sohni, A., Tan, K., Song, H.-W., Burow, D., de Rooij, D.G., Laurent, L., Hsieh, T.-C., Rabah, R., Hammoud, S.S., Vicini, E., et al. (2019). The Neonatal and Adult Human Testis Defined at the Single-Cell Level. Cell Reports 26, 1501-1517.e4. 10.1016/j.celrep.2019.01.045.

10. Shami, A.N., Zheng, X., Munyoki, S.K., Ma, Q., Manske, G.L., Green, C.D., Sukhwani, M., Orwig, K.E., Li, J.Z., and Hammoud, S.S. (2020). Single-cell RNA sequencing of human, macaque, and mouse testes uncovers conserved and divergent features of mammalian spermatogenesis. Dev Cell 54, 529-547.e12. 10.1016/j.devcel.2020.05.010.

11. Di Persio, S., Tekath, T., Siebert-Kuss, L.M., Cremers, J.-F., Wistuba, J., Li, X., Meyer zu Hörste, G., Drexler, H.C.A., Wyrwoll, M.J., Tüttelmann, F., et al. (2021). Single-cell RNA-seq unravels alterations of the human spermatogonial stem cell compartment in patients with impaired spermatogenesis. Cell Reports Medicine 2, 100395. 10.1016/j.xcrm.2021.100395.

12. Givelet, M., Firlej, V., Lassalle, B., Gille, A.S., Lapoujade, C., Holtzman, I., Jarysta, A., Haghighirad, F., Dumont, F., Jacques, S., et al. (2022). Transcriptional profiling of β-2M-SPα-6+THY1+ spermatogonial stem cells in human spermatogenesis. Stem Cell Reports 17, 936–952. 10.1016/j.stemcr.2022.02.017.

13. Izadyar, F., Wong, J., Maki, C., Pacchiarotti, J., Ramos, T., Howerton, K., Yuen, C., Greilach, S., Zhao, H.H., Chow, M., et al. (2011). Identification and characterization of repopulating spermatogonial stem cells from the adult human testis. Hum Reprod 26, 1296–1306. 10.1093/humrep/der026.

14. Tan, K., Song, H.-W., Thompson, M., Munyoki, S., Sukhwani, M., Hsieh, T.-C., Orwig, K.E., and Wilkinson, M.F. (2020). Transcriptome profiling reveals signaling conditions dictating human spermatogonia fate in vitro. Proc Natl Acad Sci U S A 117, 17832–17841. 10.1073/pnas.2000362117.

15. Shinohara, T., Orwig, K.E., Avarbock, M.R., and Brinster, R.L. (2000). Spermatogonial stem cell enrichment by multiparameter selection of mouse testis cells. Proc. Natl. Acad. Sci. U.S.A. 97, 8346–8351.

16. Zang, Z.J., Wang, J., Chen, Z., Zhang, Y., Gao, Y., Su, Z., Tuo, Y., Liao, Y., Zhang, M., Yuan, Q., et al. (2017). Transplantation of CD51+ Stem Leydig Cells: A New Strategy for the Treatment of Testosterone Deficiency. Stem Cells 35, 1222–1232. 10.1002/stem.2569.

17. Chen, P., Guan, X., Zhao, X., Chen, F., Yang, J., Wang, Y., Hu, Y., Lian, Q., and Chen, H. (2019). Characterization and differentiation of CD51+ Stem Leydig cells in adult mouse testes. Mol Cell Endocrinol 493, 110449. 10.1016/j.mce.2019.110449.

18. Willingham, S.B., Volkmer, J.-P., Gentles, A.J., Sahoo, D., Dalerba, P., Mitra, S.S., Wang, J., Contreras-Trujillo, H., Martin, R., Cohen, J.D., et al. (2012). The CD47-signal regulatory protein alpha (SIRPa) interaction is a therapeutic target for human solid tumors. Proceedings of the National Academy of Sciences 109, 6662–6667. 10.1073/pnas.1121623109.

19. Jaiswal, S., Jamieson, C.H.M., Pang, W.W., Park, C.Y., Chao, M.P., Majeti, R., Traver, D., van Rooijen, N., and Weissman, I.L. (2009). CD47 is up-regulated on circulating hematopoietic stem cells and leukemia cells to avoid phagocytosis. Cell 138, 271–285. 10.1016/j.cell.2009.05.046.

20. Miyazaki, T., Kanatsu-Shinohara, M., Ema, M., and Shinohara, T. (2023). Signal regulatory protein alpha is a conserved marker for mouse and rat spermatogonial stem cells^†^. Biol Reprod 108, 682–693. 10.1093/biolre/ioad006.

21. Mutoji, K., Singh, A., Nguyen, T., Gildersleeve, H., Kaucher, A.V., Oatley, M.J., Oatley, J.M., Velte, E.K., Geyer, C.B., Cheng, K., et al. (2016). TSPAN8 Expression Distinguishes Spermatogonial Stem Cells in the Prepubertal Mouse Testis. Biol Reprod 95. 10.1095/biolreprod.116.144220.

22. Molfetta, R., Zitti, B., Lecce, M., Milito, N.D., Stabile, H., Fionda, C., Cippitelli, M., Gismondi, A., Santoni, A., and Paolini, R. (2020). CD155: A Multi-Functional Molecule in Tumor Progression. Int J Mol Sci 21, 922. 10.3390/ijms21030922.

23. Mori, J., Nagy, Z., Di Nunzio, G., Smith, C.W., Geer, M.J., Al Ghaithi, R., van Geffen, J.P., Heising, S., Boothman, L., Tullemans, B.M.E., et al. (2018). Maintenance of murine platelet homeostasis by the kinase Csk and phosphatase CD148. Blood 131, 1122–1144. 10.1182/blood-2017-02-768077.

24. Zhu, J. (2018). Csk/CD148 and platelet SFK activation: a balancing act! Blood 131, 1042–1043. 10.1182/blood-2018-01-826438.

25. Takahashi, K., Mernaugh, R.L., Friedman, D.B., Weller, R., Tsuboi, N., Yamashita, H., Quaranta, V., and Takahashi, T. (2012). Thrombospondin-1 acts as a ligand for CD148 tyrosine phosphatase. Proc Natl Acad Sci U S A 109, 1985–1990. 10.1073/pnas.1106171109.

26. Ishii, K., Kanatsu-Shinohara, M., Toyokuni, S., and Shinohara, T. (2012). FGF2 mediates mouse spermatogonial stem cell self-renewal via upregulation of Etv5 and Bcl6b through MAP2K1 activation. Development 139, 1734–1743. 10.1242/dev.076539.

27. Masaki, K., Sakai, M., Kuroki, S., Jo, J.-I., Hoshina, K., Fujimori, Y., Oka, K., Amano, T., Yamanaka, T., Tachibana, M., et al. (2018). FGF2 Has Distinct Molecular Functions from GDNF in the Mouse Germline Niche. Stem Cell Reports 10, 1782–1792. 10.1016/j.stemcr.2018.03.016.

28. Chan, A.S., Mobley, J.L., Fields, G.B., and Shimizu, Y. (1997). CD7-mediated regulation of integrin adhesiveness on human T cells involves tyrosine phosphorylation-dependent activation of phosphatidylinositol 3-kinase. J Immunol 159, 934–942.

29. Barroca, V., Lassalle, B., Coureuil, M., Louis, J.P., Le Page, F., Testart, J., Allemand, I., Riou, L., and Fouchet, P. (2009). Mouse differentiating spermatogonia can generate germinal stem cells in vivo. Nat. Cell Biol. 11, 190–196. 10.1038/ncb1826.

30. Futamura, K., Sekino, M., Hata, A., Ikebuchi, R., Nakanishi, Y., Egawa, G., Kabashima, K., Watanabe, T., Furuki, M., and Tomura, M. (2015). Novel full-spectral flow cytometry with multiple spectrally-adjacent fluorescent proteins and fluorochromes and visualization of in vivo cellular movement. Cytometry A 87, 830–842. 10.1002/cyto.a.22725.

31. Van Gassen, S., Callebaut, B., Van Helden, M.J., Lambrecht, B.N., Demeester, P., Dhaene, T., and Saeys, Y. (2015). FlowSOM: Using self-organizing maps for visualization and interpretation of cytometry data. Cytometry A 87, 636–645. 10.1002/cyto.a.22625.

32. Bendall, S.C., Davis, K.L., Amir, E.D., Tadmor, M.D., Simonds, E.F., Chen, T.J., Shenfeld, D.K., Nolan, G.P., and Pe’er, D. (2014). Single-Cell Trajectory Detection Uncovers Progression and Regulatory Coordination in Human B Cell Development. Cell 157, 714–725. 10.1016/j.cell.2014.04.005.

