## Supplemental Figures for "Characterization and hierarchy of the spermatogonial stem cell compartment in human spermatogenesis by spectral cytometry using a 16-colors panel"

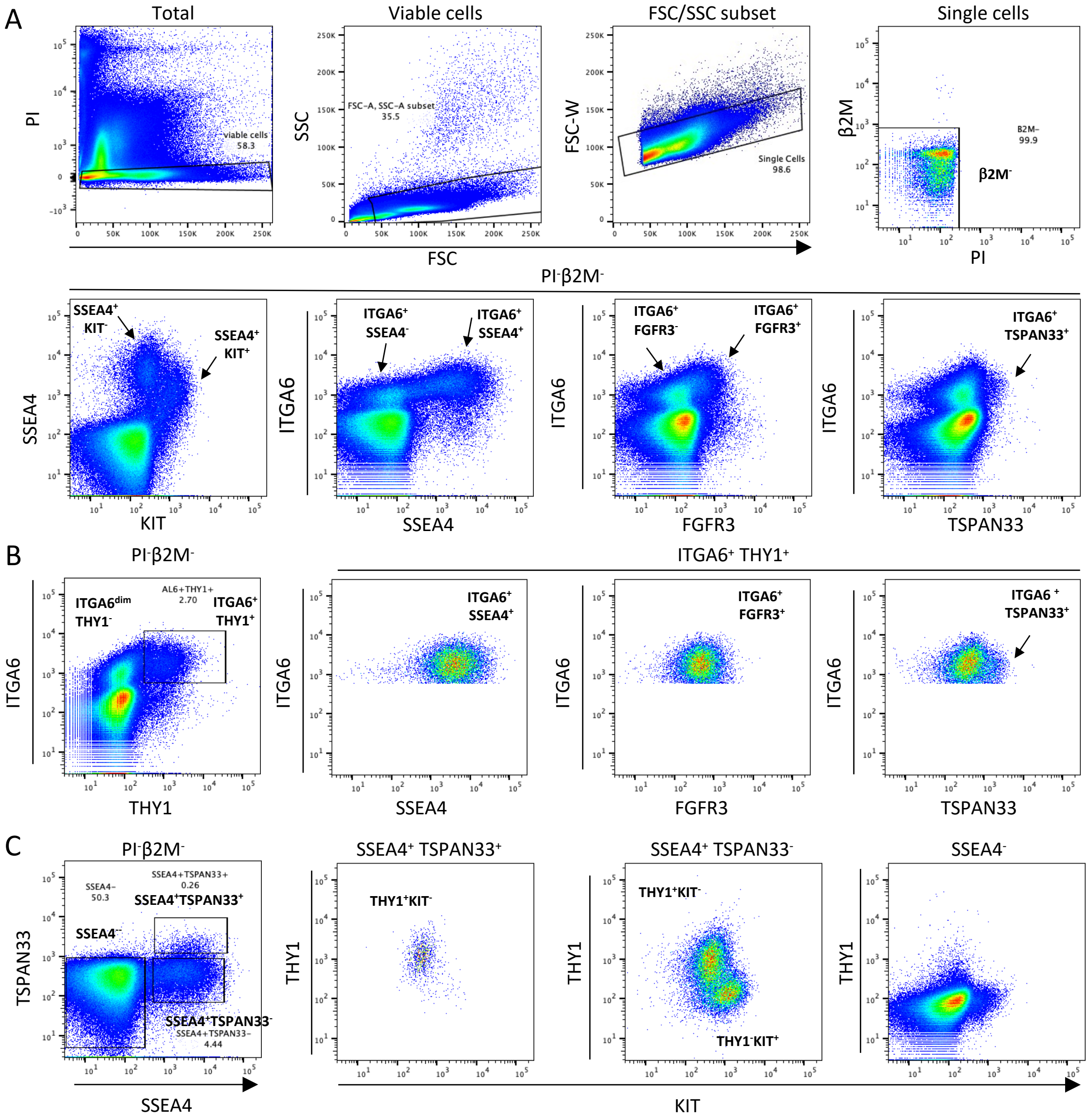


Figure S1: Characterization of SSCs and spermatogonial progenitors using 8-C fluorescence panel.

(A) Representative flow cytometry plots of the 8-C panel analysis on the human spermatogonial progenitors according to forward (FSC) and side scatter (SSC), β2 microglobulin, ITGA6, SSEA4, KIT, THY1, FGFR3, and TSPAN33 parameters. Viable propidium iodide (PI)-negative cells were analyzed using forward and side scatter to remove sperm from further analyses, and single cells were selected on FSCA-FSCW. Population frequencies are shown in the graph. Side population marker has been omitted from this 8-C panel in order to offer new possibilities of fluorochromes to be used on the violet or UV lasers, the use of Vybrant or Hoechst 33342 DNA dye preventing the detection of a higher number of markers. (B) The $\beta 2M^{-}ITGA6^{+}THY1^{+}$ population contains the spermatogonia expressing SSEA4, FGFR3, and the fraction of cells expressing TSPAN33 (C) Flow cytometric cytogram of TSPAN33 and SSEA4 markers in the $\beta 2M^{-}$ germinal population, and THY1 and KIT expression in $SSEA4^{+}TSPAN33^{+}$, $SSEA4^{+}TSPAN33^{-}$, $SSEA4^{-}TSPAN33^{-}$ germinal populations. $SSEA4^{+}TSPAN33^{-}$ cells are a mixture of $THY1^{+}KIT^{-}$ cells and $THY1^{-}KIT^{+}$ cells.

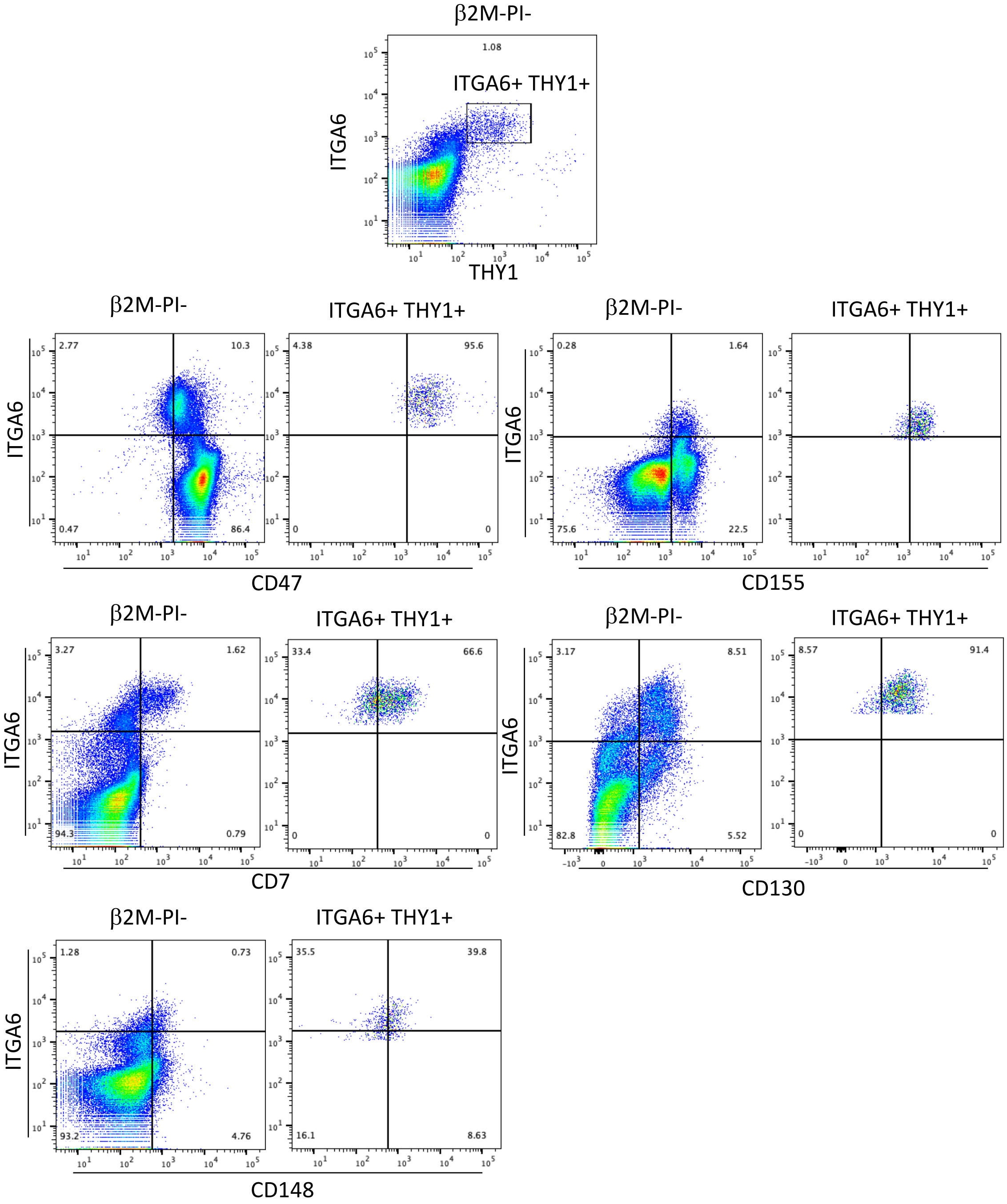


Figure S2: Flow cytometric analysis of the Expression of CD7, CD47, CD155, CD148, and CD130 in the whole β2M$^{-}$ germinal population and in the β2M$^{-}$ITGA6$^{+}$THY1$^{+}$ population. CD7, CD47, CD130, CD155, and CD148 are new markers of β2M$^{-}$ITGA6$^{+}$THY1$^{+}$ spermatogonia. For the screening, we studied the expression in the β2M$^{-}$ITGA6$^{+}$THY1$^{+}$ population of 17 candidate markers (CD7, TSPAN8, CD47, CD77, CD24, CD155, CD184, CD148, CD51/CD61, CD130, EPHB2, CD115, CD146, CD226, CD304/NRP1, CD334/FGFR4, and CD75). Apart from CD7, CD47, CD130, CD155, and CD148, the other markers were negative for the antibody clones we used for screening.

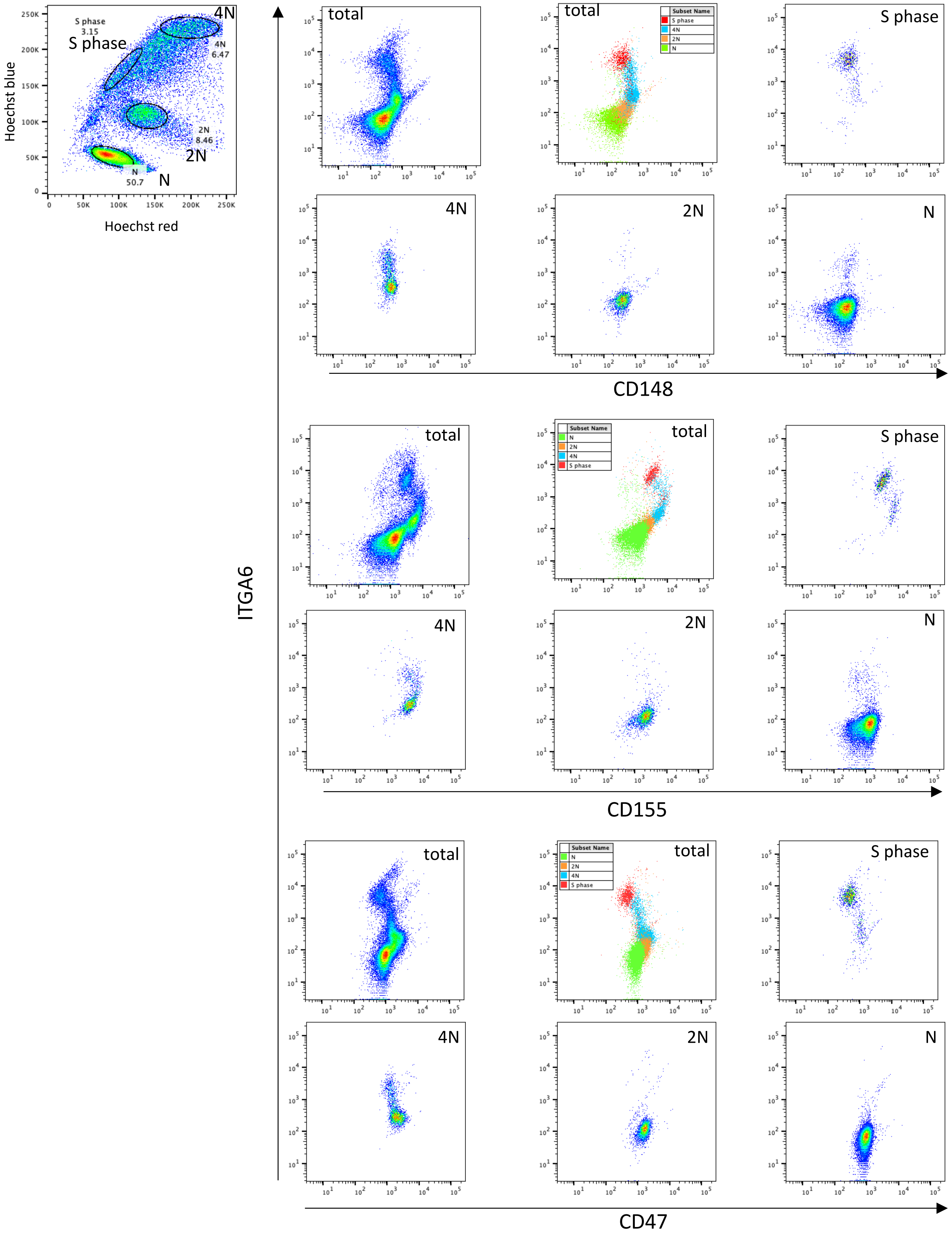


<u>Figure S3:</u> Assignment of meiotic and post meiotic populations according to the expression of ITGA6, CD148, CD155, and C47. Late differentiating spermatogonia "S-phase" ("S-phase"), spermatocyte (4N), spermatocyte II (2N), and spermatids (N) germinal populations were defined according to their DNA content (Hoechst 33342 fluorescence) as previously described[12].

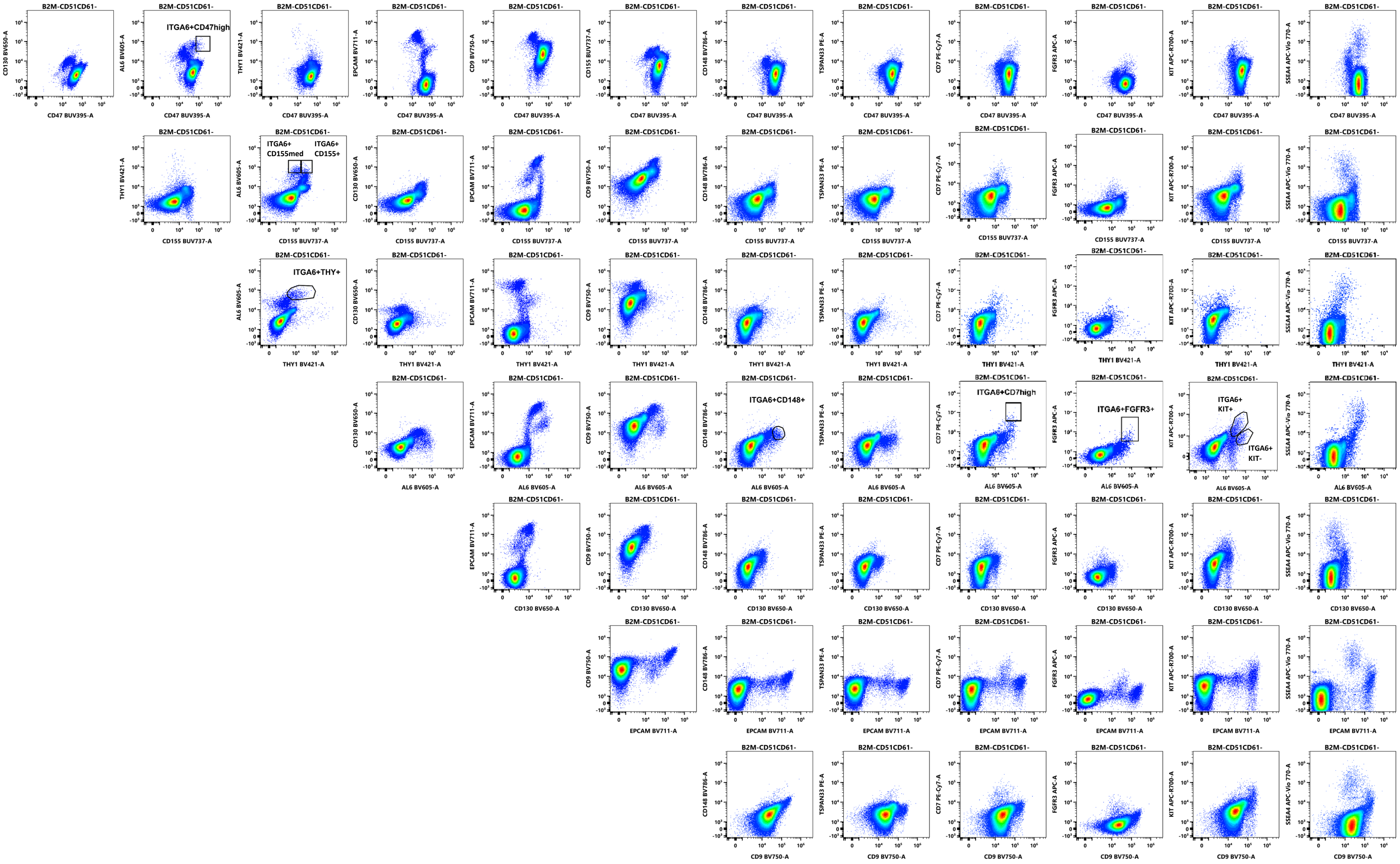


Figure S4: Representative flow cytometry plots of the 13 germinal markers gated on $\beta$2M$^-$CD51/CD61$^-$ cells. NxN plot displaying every parameter versus every other parameter, gates of specific germinal populations are indicated on plots. (ITGA6 is named AL6 in cytogram axis)

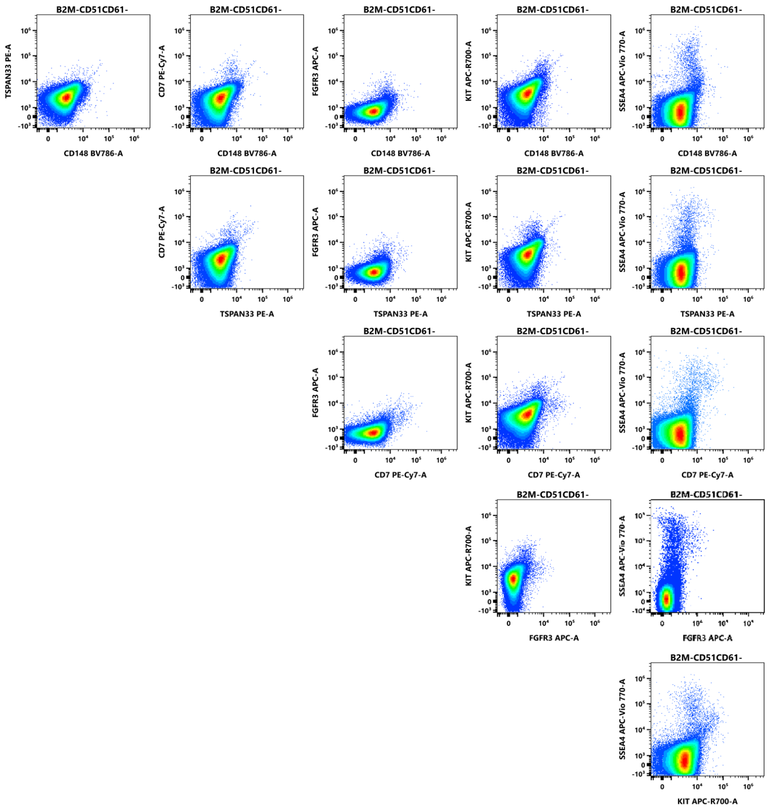


<u>Figure S4 (continued):</u> Representative flow cytometry plots of the 13 germinal markers gated on $\beta 2M^{-}CD51/CD61^{-}$ cells . NxN plot displaying every parameter versus every other parameter, gates of specific germinal populations are indicated on plots. (ITGA6 is named AL6 in cytogram axis)

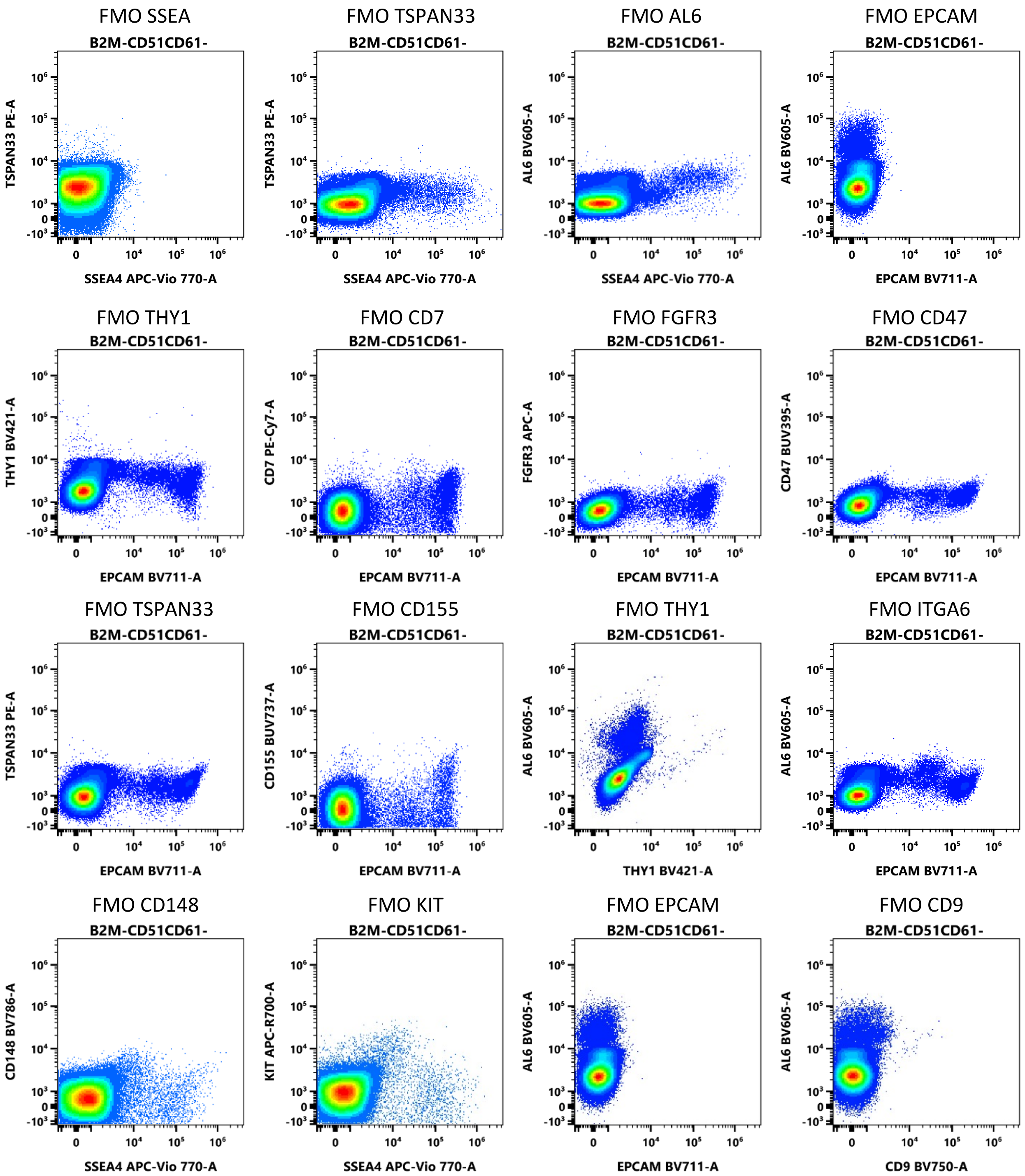


<u>Figure S5</u>: Fluorescence minus one (FMO) relevant controls for the 13 germinal markers used for flow cytometry profiling of adult testicular cell samples. (ITGA6 is noted AL6 in cytogram axis)

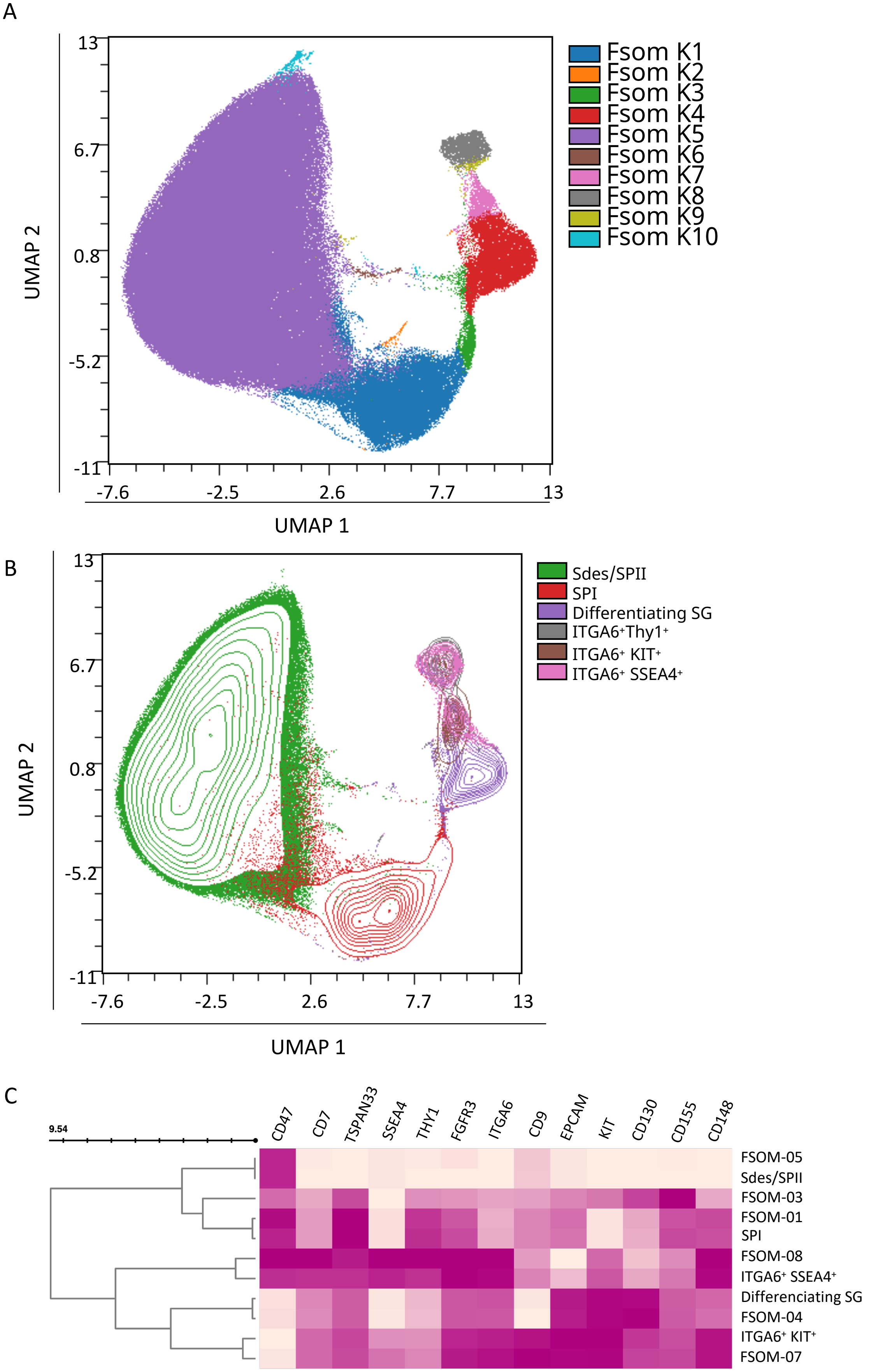


Figure S6: Unsupervised FlowSOM analysis of the multi-dimensional 16-C panel dataset on human germinal cells. (A) High-dimensional UMAP analysis of $\beta$2M/CD51/CD61$^-$ cells with overlay of the 10 FlowSOM clusters and (B) of the testicular subsets defined in Fig. 1, Figure S3 and Figure S4. (C) Heatmap of the expression pattern of individual spermatogonial markers in the different germinal subsets and FlowSOM clusters.

A

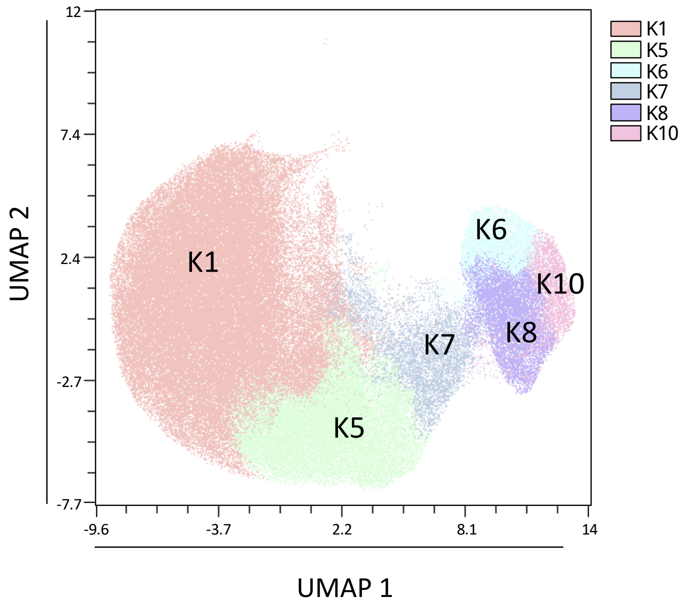


B

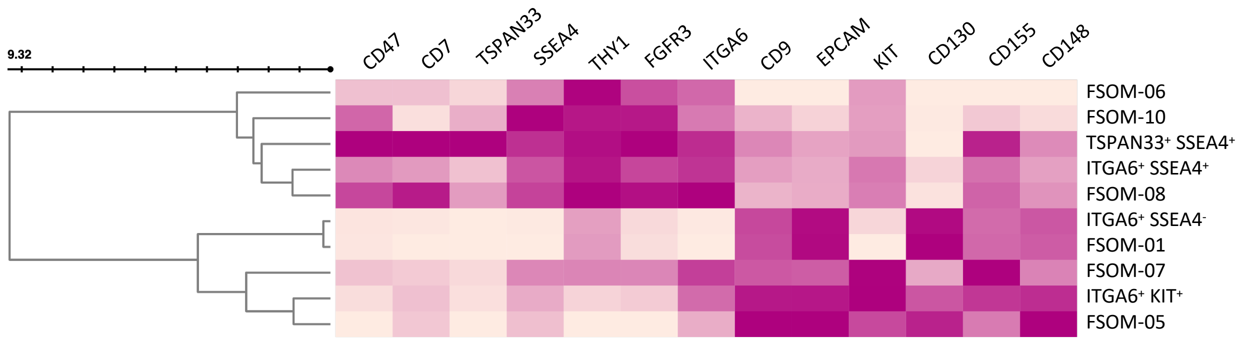


<u>Figure S7:</u> Unsupervised FlowSOM analysis of the 16-C panel dataset on ITGA6$^+$ spermatogonia. (A) High-dimensional UMAP analysis of β2M/CD51/CD61$^-$ITGA6$^+$ cells with overlay of the 6 major FlowSOM clusters. (B) Heatmap of the expression pattern of individual spermatogonial markers in the different testicular subsets defined in Fig. 1 and Figure S4, and in the FlowSOM clusters.

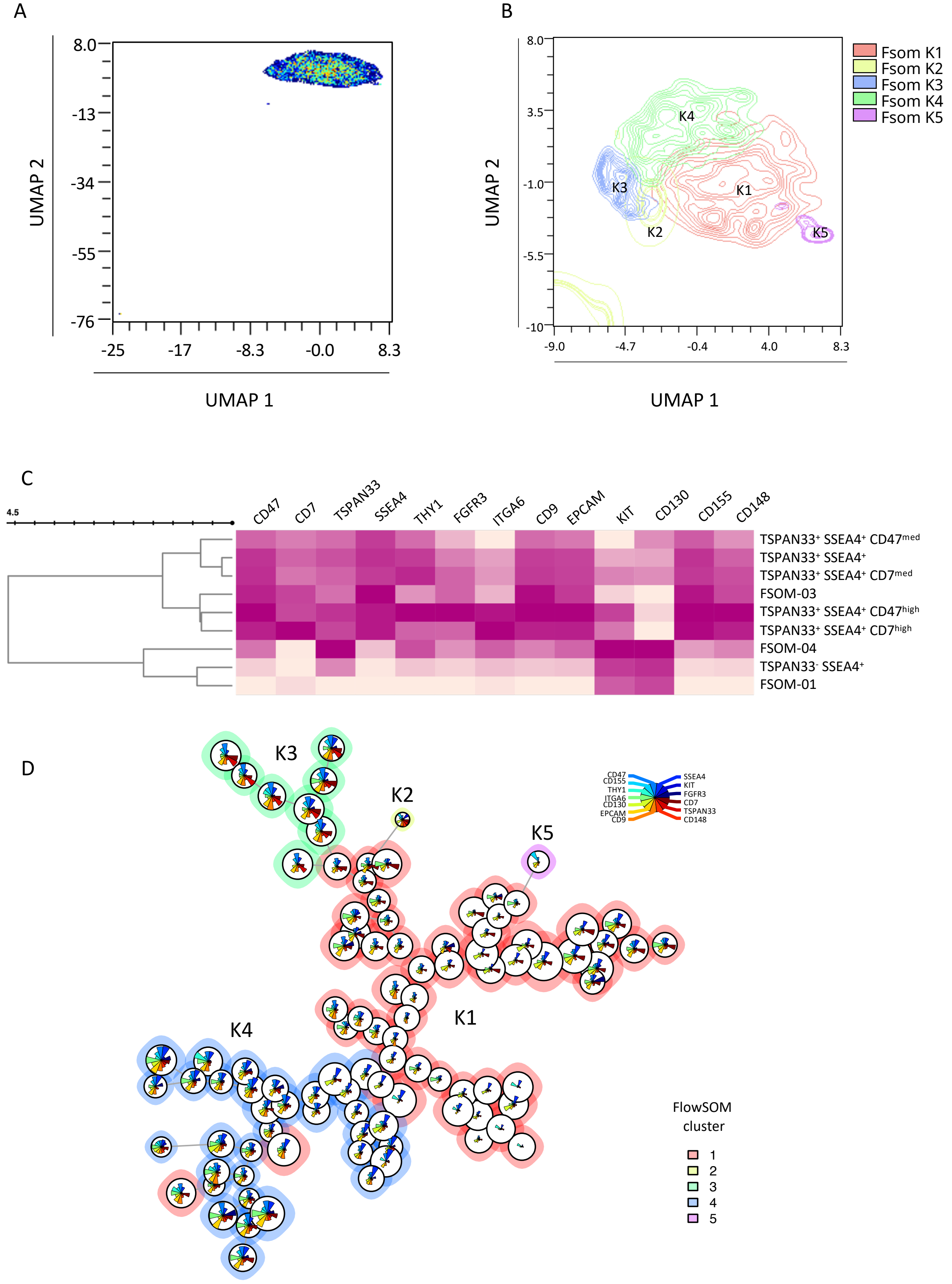


Figure S8: Unsupervised FlowSOM analysis of the 16-C panel dataset on ITGA$^{+}$THY1$^{+}$ spermatogonial population. (A) High-dimensional UMAP analysis of β2M/CD51/CD61$^{-}$ ITGA6$^{+}$THY1$^{+}$ cells (B) High-dimensional UMAP analysis of β2M/CD51/CD61$^{-}$ITGA6$^{+}$THY1$^{+}$ cells with overlay of the 5 FlowSOM clusters (zoom on the main populations, see Figure 6A). (C) Heatmap of the expression pattern of individual spermatogonial markers in the different testicular subsets defined in Figure 1, Figure S4, and in the FlowSOM clusters. (D) FlowSOM spermatogonial subsets are indicated on the FlowSOM tree representation.

| Primers | Sequences |
|---|---|
| UTF1 F | 5'-CGGCTCCCAGCGAACCAG-3' |
| UTF1 R | 5'-GACGGGCTGAAGCGGAGC-3' |
| C19orf84 F | 5'-AGATGGAACAACCAAAGGACG-3' |
| C19orf84 R | 5'-GTTCAGGACAAGGGTGGAG-3' |
| LPPR3 F | 5'-CTTCTGCCCTGCTTCTACTTCG-3' |
| LPPR3 R | 5'-CATAGCACTGGAAGCCCACC-3' |
| PIWIL4 F | 5'-CATCAAGTTCTCCCGTGTGC-3' |
| PIWIL4 R | 5'-GACACAGAAATGGCAAACCC-3' |
| FGFR3 F | 5'-CCGAGCGGATGGACAAGAAG-3' |
| FGFR3 R | 5'-GACCAGGCTCCACTGCTGAT-3' |
| GAPDH F | 5'-GAAGGTGAAGGTCGGAGTCA-3' |
| GAPDH R | 5'-TGGACTCCACGACGTACTCA-3' |

Table S1 : list of primers
